# Analysis of noise mechanisms in cell size control

**DOI:** 10.1101/080465

**Authors:** Saurabh Modi, Cesar A. Vargas-Garcia, Khem Raj Ghusinga, Abhyudai Singh

**Author notes:** Corresponding Author: Abhyudai Singh,.

## Abstract

At the single-cell level, noise features in multiple ways through the inherent stochasticity of biomolecular processes, random partitioning of resources at division, and fluctuations in cellular growth rates. How these diverse noise mechanisms combine to drive variations in cell size within an isoclonal population is not well understood. To address this problem, we systematically investigate the contributions of different noise sources in well-known paradigms of cell-size control, such as the adder (division occurs after adding a fixed size from birth) and the sizer (division occurs upon reaching a size threshold). Analysis reveals that variance in cell size is most sensitive to errors in partitioning of volume among daughter cells, and not surprisingly, this process is well regulated among microbes. Moreover, depending on the dominant noise mechanism, different size control strategies (or a combination of them) provide efficient buffering of intercellular size variations. We further explore mixer models of size control, where a timer phase precedes/follows an adder, as has been proposed in *Caulobacter crescentus*. While mixing a timer with an adder can sometimes attenuate size variations, it invariably leads to higher-order moments growing unboundedly over time. This results in the cell size following a power-law distribution with an exponent that is inversely dependent on the noise in the timer phase. Consistent with theory, we find evidence of power-law statistics in the tail of *C. crescentus* cell-size distribution, but there is a huge discrepancy in the power-law exponent as estimated from data and theory. However, the discrepancy is removed after data reveals that the size added by individual newborns from birth to division itself exhibits power-law statistics. Taken together, this study provides key insights into the role of noise mechanisms in size homeostasis, and suggests an inextricable link between timer-based models of size control and heavy-tailed cell size distributions.

## Introduction

Unicellular organisms employ diverse control strategies to maintain size homeostasis i.e., to ensure that they do not become abnormally large (or small) [1–5]. It is well known that cells within an isoclonal population, which presumably follow identical size-control strategies, can exhibit significant intercellular differences in size [6–10]. Here we systematically explore how such stochastic variation in cell size is impacted by various underlying noise sources, such as:

- Noise in partitioning of volume among daughter cells during mitosis and cytokinesis [11–13].
- Random fluctuations in the cell growth-rate that potentially have memory over multiple generations [14–18].
- Stochasticity in the biomolecular processes associated with the cell cycle that generate randomness in the timing of cell-division [19–21].

A key question of interest is whether stochastic variation in cell size is more sensitive to some noise sources over others. Moreover, how does this sensitivity to noise mechanisms change across size-control strategies.

We investigate the above questions in the context of the recently uncovered “adder strategy” for size homeostasis. As per this strategy, division is triggered after newborn cells add (on average) a constant size to their size at birth [22–25]. Assuming exponential growth in cell size over time, the adder implies that larger newborns divide earlier (i.e., the constant size is accumulated in shorter time) than smaller newborns. The generality of this strategy can be underscored from that fact it has been reported in many microbial species, such as, *E. coli* [24], *B. subtilis* [24], *P. aeruginosa* [26], and *D. Hildenborough* [27]. We begin by describing the stochastic formulation of the adder model that encompasses different noise sources consistent with findings of recent single-cell studies [14, 22]. Later on, this model is expanded to “the generalized adder”, that encapsulates both the adder, and the sizer (division occurs upon reaching a size threshold) paradigms of size control [24, 28].

## Stochastic formulation of cell size control

Consider tracking an individual cell undergoing cycles of exponential growth and division. Let *V*_*n*_ denote the size of the (newborn) cell at the start of the *n*^*th*^ cell cycle. In between successive division events, size increases exponentially with rate 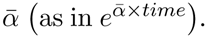. Then, as per the adder strategy,

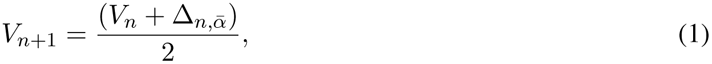

where the random variable 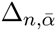 denotes the size added to *V*_*n*_ just before the mother cell divides into two equally-sized daughters. To adopt a form for 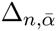, we consider two key experimental insights from *E. coli*:

- For a given 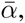 the histograms of the added size corresponding to different newborn sizes collapse on top of each other [14]. This implies that not only the mean, but the entire distribution of 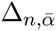 is invariant of *V_n_*.
- While the mean size added 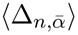 depends on 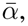 the distribution of size added normalized by its mean 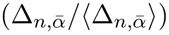 becomes invariant of it [14]. Thus, varying growth conditions essentially rescales the distribution of size added by its corresponding mean.

These findings motivate the following form for the added size

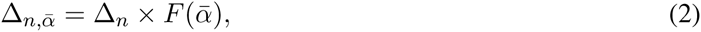

where ∆_*n*_ is an independent and identically distributed (iid) random variable following a size-independent distribution. ∆_*n*_ is assumed to have the following mean and coefficient of variation squared

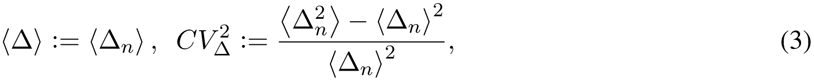

respectively, where the symbol ‹ › denotes the expected value. Throughout the manuscript, the subscript *n* is dropped for describing the statistical moments of iid random variable like ∆_*n*_. Randomness in ∆_*n*_ essentially encompasses noise inherent in the processes of cell cycle, and timing of cell division. In light of this, we refer to 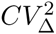 as the extent of *cell-division noise*. *F* in (2) represents a non-decreasing function of the growth rate, and empirical data suggests that it can have a linear 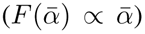 [26] or exponential 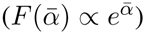 form [14]. Next, we incorporate another noise source that critically impacts size fluctuations - errors incurred in the partitioning of the mother cell volume between two daughters.

## Incorporating noise in size partitioning

Stochasticity in the partitioning process is accounted by modifying (2) to

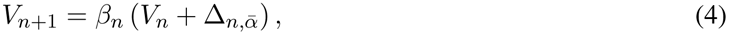

where *β*_n_ *∈* (0, 1) is an iid random variable with mean *‹β›* = 1/2 and coefficient of variation squared 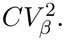. We refer to 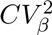 as the magnitude of *partitioning error*. Given the restriction on the support of *β*_*n*_, it is conveniently modeled through a beta distribution with appropriately chosen parameters so as to match *‹β›* and 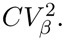 Note that this framework applies for both symmetric and asymmetric partitioning. If after cell division, the newborn size *V*_n+1_ is found by randomly selecting one of the daughter cells, then *‹β›* = 1/2 even for asymmetric partitioning, albeit with a much higher 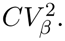 The partitioning error 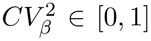 is always bounded with the maximum value occurring in the case of extreme asymmetric partitioning

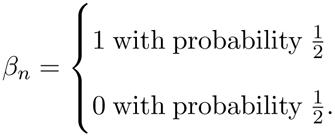

Scenarios where *V*_n+1_ is found by selecting daughter cells in a biased fashion (for example, the large cell is always chosen), can be incorporated by allowing *‹β›* ≠ 1/2

## Incorporating growth-rate fluctuations

Having introduced two different mechanisms for driving stochastic variations in cell size (cell-division noise and partitioning errors), we consider the third and final noise mechanism – fluctuations in the growth rate that potentially arise from noisy expression of metabolic enzymes [15]. Let *α_n_* denote the growth rate of a newborn cell at the start of the *n^th^* cell cycle. The time evolution of *α_n_* is modeled through the following discrete-time autoregressive model

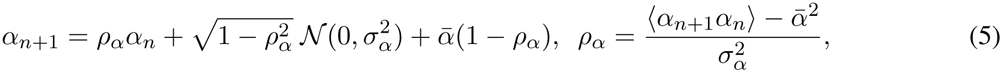

where *ρ*_*α*_ *∈* [0, 1] denotes the correlation of growth rates across cell cycles, 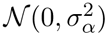 represents iid zero-mean random variables with variance 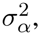, and 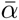 is the mean growth rate. A key quantity of interest is the coefficient of variation of 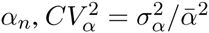 that measures the magnitude of fluctuations in the growth rate.

A natural question that arises at this point is how to connect *α*_*n*_ to the size added by a single newborn cell before division. In many growth conditions, the size added by single *E. coli* newborns shows little or no dependence on *α*_*n*_ [22], in which case growth-rate fluctuations can be ignored. However, single-cell correlations between the size added and *α*_*n*_ have been reported for some growth mediums [22]. To capture this effect we take a phenomenological approach and modify 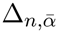 in (4) to

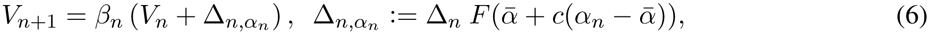

where *c ∈* [0, 1]. Here, *c* = 0 corresponds to the scenario where the size added, and the growth rate, are connected via their population averages 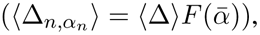 but are independent at the single-cell level. In contrast, *c* = 1 represents strong coupling between them at both the population and single-cell levels. In the limit of small fluctuations in *α*_*n*_

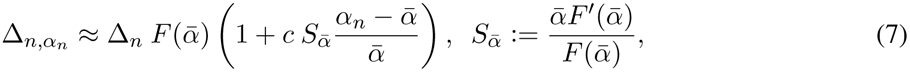

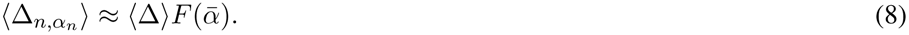

where 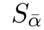 is the log sensitivity of the function *F* to the growth rate, and is given by 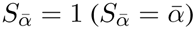 when *F* takes a linear (exponential) form [14]. Note that if fluctuations in *α*_*n*_ are uncorrelated across cell cycles (*ρ*_*α*_ = 0), then ∆_*n,α*_*n*__ is completely independent of the newborn cell size *V*_*n*_. However, weak dependencies between ∆_*n,α*_*n*__ and *V*_*n*_ may arise for *ρ*_α_ *>* 0 and *c >* 0.

## The generalized adder

To enhance the repertoire of size control strategies, the discrete-time model (6) is expanded to

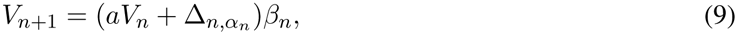

where *a* ∈ [0; 1], and *aV*_*n*_+ ∆_*n,α*_*n*__ is the cell size just before division. This model has often been referred to in literature as the generalized adder [28]. While *a* = 1 corresponds to the adder, a = 0 is the well-known sizer strategy,

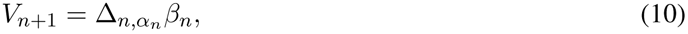

where division occurs when the cell reaches a size threshold ∆_*n,α*_*n*__ defined in (7) [16, 29, 30]. Note that the biological interpretation of ∆_*n,α*_*n*__ changes with *a* – it is the size added (size threshold) for division to occur as per the adder (sizer) control strategy that is realized when *a* = 1 (*a* = 0). Values of *a* between 0 and 1 could denote imperfect implementations of the adder and sizer, or represent combinatorial size control using both strategies [31]. In summary, the overall model describing the stochastic dynamics of newborn cell size is given by (5), (7) and (9). Among the three noise sources impacting cell size, cell-division noise and growth-rate fluctuations are additive noise mechanisms implemented via ∆_*n,α*_*n*__, while *β_n_* is a multiplicative noise mechanism. Our goal is to quantify the steady-state moments of *V_n_*, and investigate how they are impacted by the magnitude of noise sources - 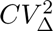 (cell-division noise), 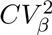 (partitioning errors) and 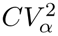 (growth-rate fluctuations). To guide the reader, a description of the various symbols used in the paper are provided in Table 1.

**Table 1:**
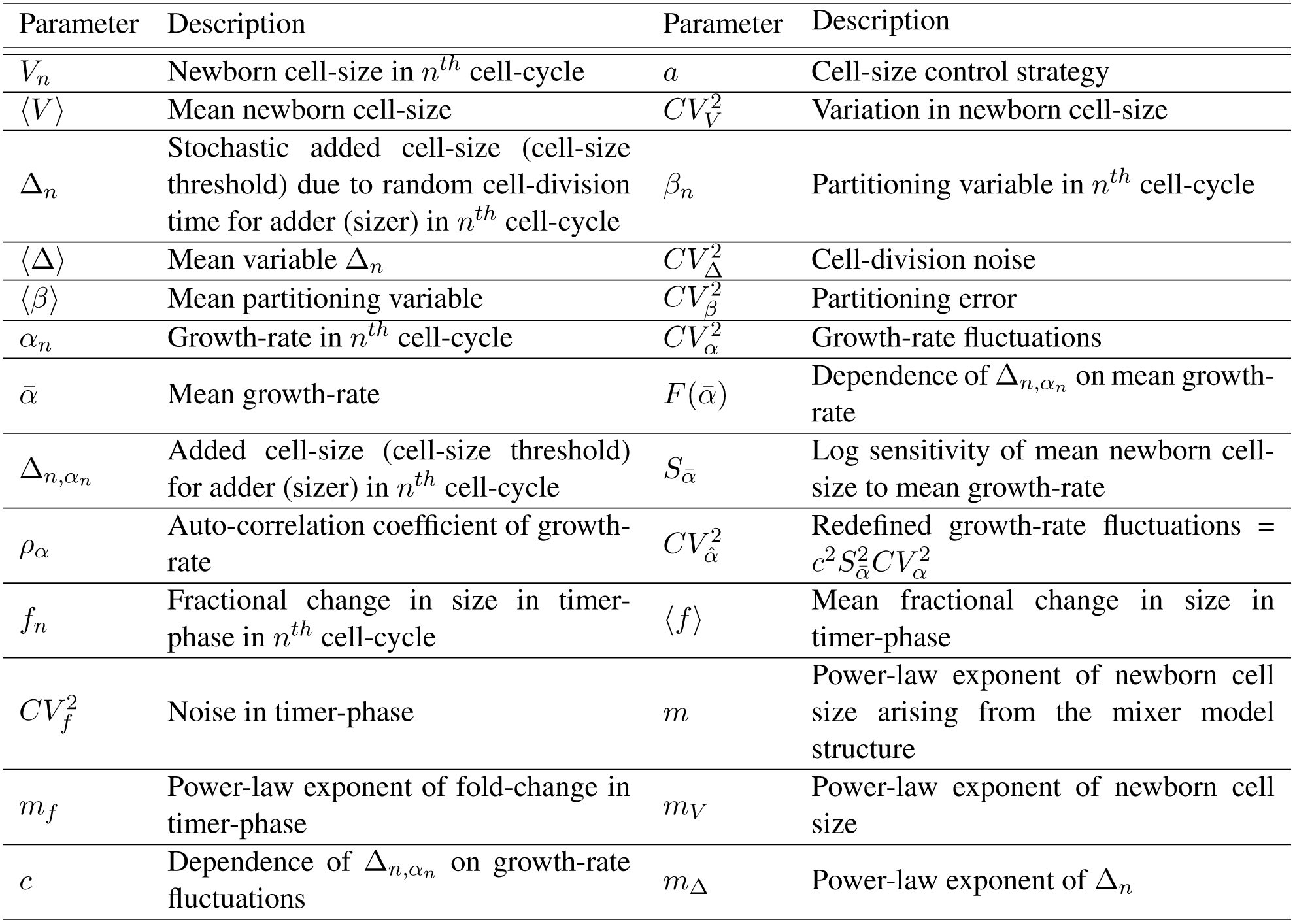
Summary of notation used in the paper

## Analysis of mean and variation in newborn cell size

Let the steady-state mean and noise (as quantified by the coefficient of variation squared) levels of the newborn cell-size be denoted by

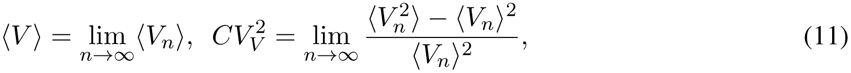

respectively. Using the fact that *β*_*n*_ is drawn independently of ∆_*n,α*_*n*__ and *V*_*n*_, it is straightforward to show from (9) that

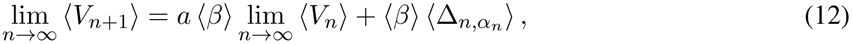

which using 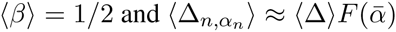 reduces to

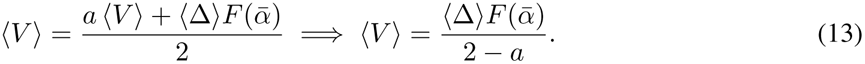

Thus, the mean newborn size increases with growth rate as per the function *F*. Moreover, the adder yields larger cells on average, than the sizer, assuming same ‹∆› for both strategies.

A similar analysis for the second-order moment yields the following noise level (see Section S1 in SI)

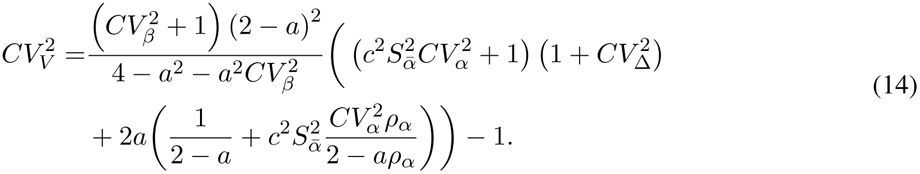

Note that in light of (13), 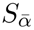 defined in (7) can now be interpreted as the log sensitivity of the mean newborn cell size to the average growth rate 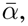. To gauge the effect of different noise mechanisms, we plot 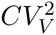 as a function of individual noise magnitudes assuming other noise sources are absent. Results show that for 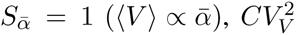increases most rapidly with 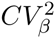 (Fig. 1). Thus, effective regulation of partitioning errors is a key ingredient for buffering stochastic variations in size. Perhaps not surprisingly, (14) further shows that 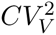 increases as

**Figure 1:**
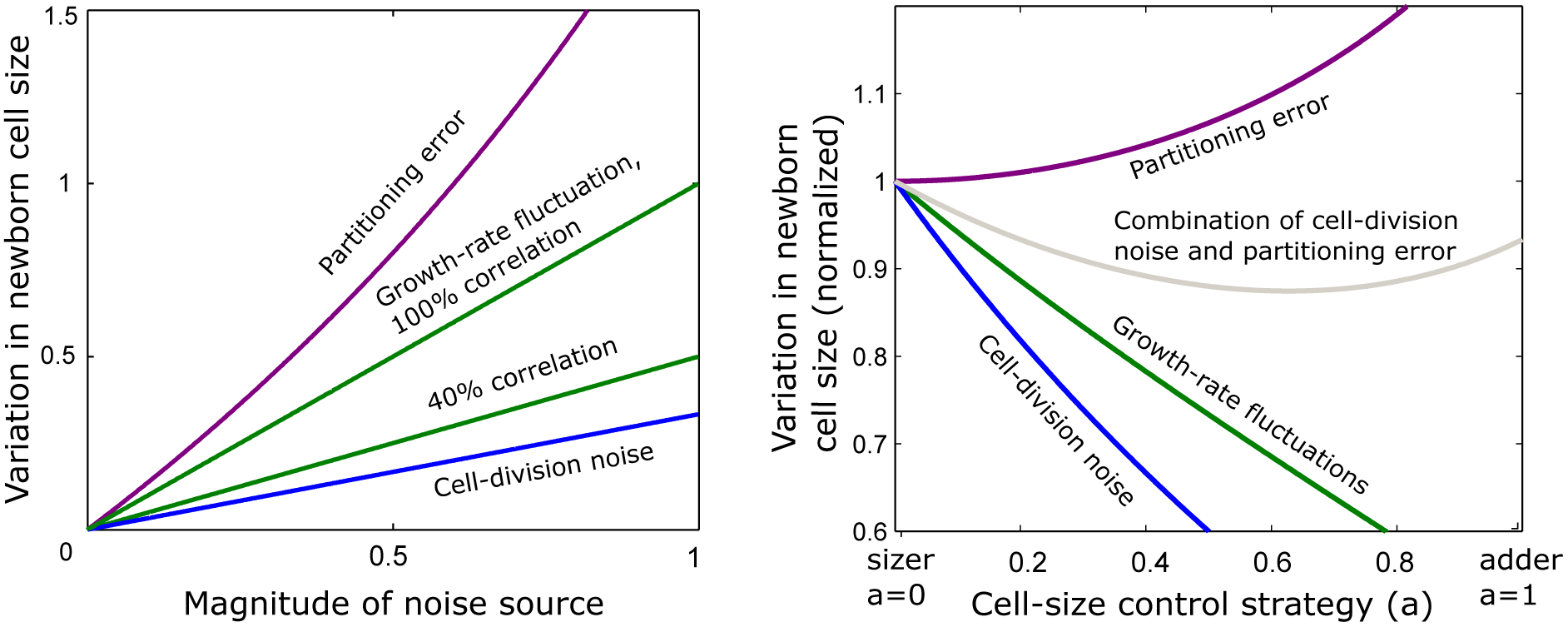
Stochastic variations in newborn cell size are most sensitive to partitioning errors. *Left*: The extent of size variations 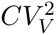 given by (14) are plotted as a function of partitioning error 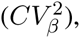, cell-βdivision noise 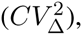, and growth-rate fluctuations 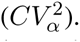. The latter plot of 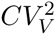 against 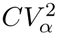 is for growth rate correlations *ρ*_*α*_ = 0.4 and 1. For each line, other noise sources are set to zero. Results show that size variations arising from partitioning errors dominate the contributions of other noise sources. Parameters taken as *a* = 1 (adder) and 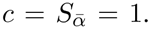. *Right*: The noise contributions in (16) are plotted as a function of *a*. Each contribution is normalized by its value for *a* = 0. The noise contribution from growth-rate fluctuations is plotted for *ρ_α_* = 0.4. The combined contribution from cell-division noise and partitioning errors for 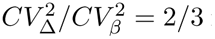 is minimized at an optimal value of *a*.

1. Fluctuations in the growth rate become correlated across generations (i.e., increasing *ρ*_*α*_)
2. ‹*V*› becomes more sensitive to 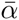 (i.e., increasing 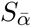)
3. The size added ∆_*n,α*_*n*__ becomes correlated with *α*_*n*_ at the level of individual cells (i.e., increasing *c*).

Since 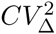 always appears in (14) with 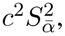, for the remainder of the manuscript we redefine the extent of

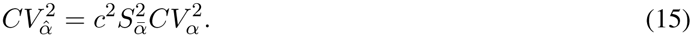

## Analysis of newborn cell size for low source noises

Stochastic variations in cell size are generally well regulated such that 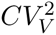 values are significantly smaller than one. For example, size variation across a clonal population of *E. coli* cells has been reported to be less than 35% under different growth conditions, implying 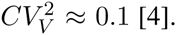 [4]. Exploiting small noise magnitudes 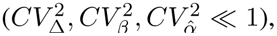 linearization of (14) allows decomposition of 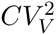 into components representing contributions of individual noise mechanisms

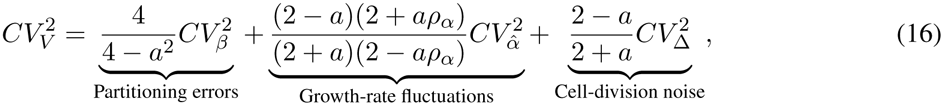

where the linear coefficients can be interpreted as the sensitivity of 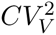 to different noise sources. For example, 4/(4 *− a*^2^) is the sensitivity of 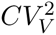 to partitioning errors, and lower sensitivities correspond to more effective control of size variations. For *a ∈* (0, 1], the sensitivities in (16) always satisfy

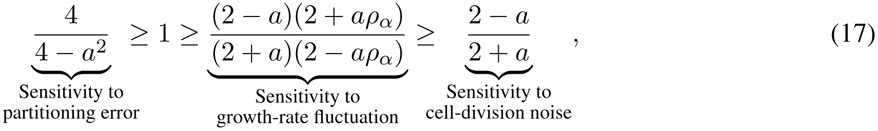

implying that size variations are most sensitive to partitioning errors consistent with the findings of Fig.1 Note that the sensitivity of 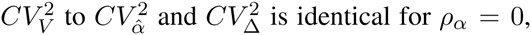 with the former increasing as growth-rate fluctuations become more correlated across cell cycles. With increasing *a*, the sensitivity of 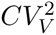 to partitioning errors increases, however, its sensitivity to noise mechanisms in ∆_*n,α*_*n*__ (i.e., cell-division noise and growth-rate fluctuations) decreases (Fig. 1). This effect is exemplified by

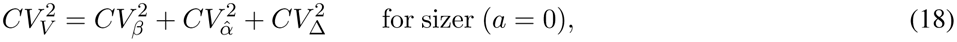

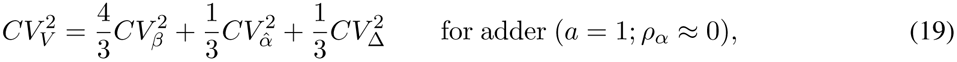

where the adder has higher (lower) sensitivity to partitioning errors (cell-division noise and growth-rate fluctuations) as compared to the sizer. Thus, while the adder provides more effective noise buffering against cell-division noise and growth-rate fluctuations, the sizer is a better strategy against partitioning errors. Intuitively, in sizer-based control (10), the newborn size is only affected by noise in the previous cell cycle. In contrast, size is affected by noise in *all* previous cell cycles in the adder. This dependence on history in the adder is a double-edged sword as it allows effective averaging of additive noise mechanisms like ∆_*n,α*_*n*__ across multiple cell cycles, but this comes at the cost of amplifying multiplicative noise mechanisms like *β*_*n*_. Given this trade-off, when both partitioning and cell-division noise are present at comparable levels, 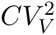 values are minimized by a combination of adder and sizer, i.e., at intermediate values of *a* (Fig. 1).

In summary, simple stochastic models based on recurrence equations provide critical insights into contributions of alternative noise mechanisms to size fluctuations, and what control strategies provides the best buffering of fluctuations.

## Mixer models for size control

In many organisms like *Caulobacter crescentus*, size control is exerted in only a part of the cell cycle [32, 33]. More specifically, the cell cycle can be divided into two phases - a timer phase where the cell grows exponentially for a size-independent duration of time. In essence, this phase represents time required to complete a certain cell-cycle step (such as, genome replication) irrespective of size. The timer is followed by a generalized-adder phase that regulates size as per (9). We refer to such biphasic size control as mixer models, and investigate how the duration and noise in the timer phase alters the noise contributions in (16).

Consider a newborn cell that grows in the timer phase from *V*_*n*_ to (1 + *f*_*n*_)*V*_*n*_, where *f*_*n*_ > 0 is an iid random variable drawn from an arbitrary positively-valued distribution. Based on this formulation, 1 + ‹*f*› is the average fold-change in cell size in the timer phase and we assume ‹*f*› < 1 (size increase is less than two-fold). Furthermore, the noise in timer phase is quantified by 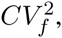 the coefficient of variation squared of *f*_*n*_. A generalized adder controlling size from the end of timer phase to cell division, yields the following mixer model

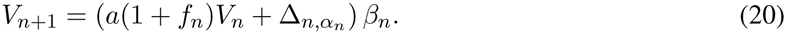

While (20) assumes that the timer precedes the generalized adder, the opposite scenario of timer following the generalized adder results in

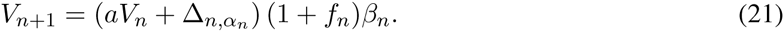

Note that both models converge to (9) in the limit of *f*_*n*_ = 0. The average newborn sizes corresponding to (20) and (21) are given by

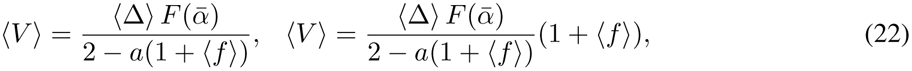

respectively, and shows that the former case where size regulation occurs at the end of cell cycle (i.e., timer precedes the generalized adder) results in smaller cells. Next, we quantify and investigate cell-size variations in mixer models.

## Noise analysis of mixer models

Analysis of (20) (timer precedes the generalized adder) yields the following convoluted formula for the steady-state coefficient of variation of *V_n_* (SI section S2)

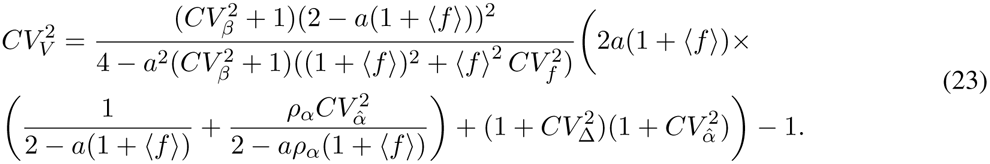

The corresponding formula for the opposite case (21) (timer follows the generalized adder) is

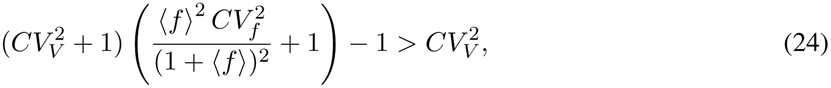

and reveals elevated noise levels in *V*_*n*_ compared to (23). This is expected since in (21) the timer-phase noise affects both terms *aV*_*n*_ and ∆_*n,α*_*n*__, while in (20) only one term is affected.

Assuming 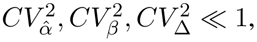, the variation in newborn size 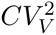 can be decomposed into individual noise contributions

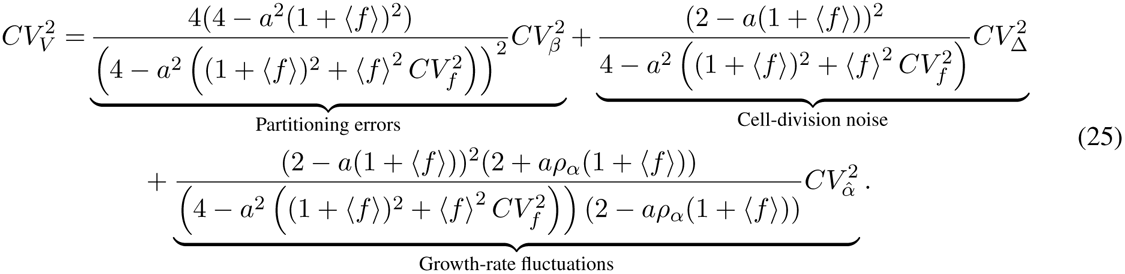

To study the sensitivity of 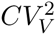 to different noise sources, the linear coefficients are plotted as a function of ‹*f*› for two different levels of the timer-phase noise 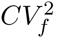 (Fig. 2). Our analysis shows that the sensitivity of 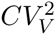 to partitioning errors increases dramatically with the inclusion of a timer phase, and the sensitivity blows up for finite values of ‹*f*›. In contrast, sensitivity to 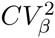 and 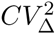 first decreases and then increases exhibiting a U-shape profile, with the dip in the profile being more pronounced for low levels of timer-phase noise (Fig. 2). An important implication of this is that the inclusion of a precise timer phase in the generalized adder can reduce size variations when partitioning errors are negligible.

**Figure 2:**
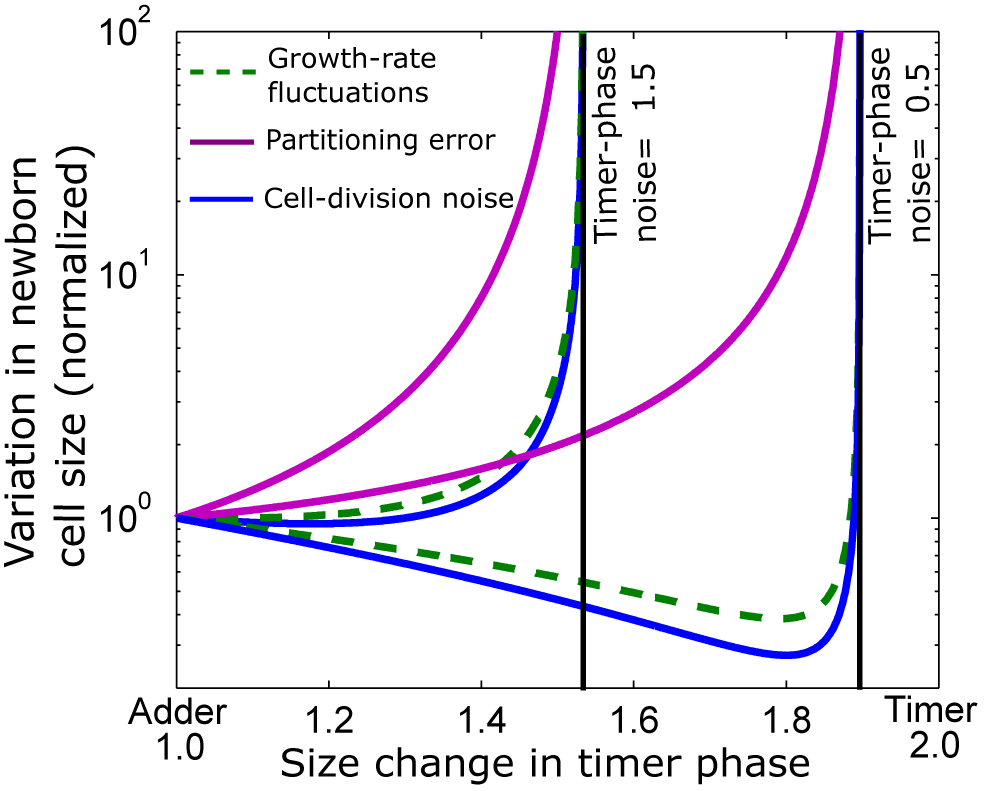
The contributions of different noise mechanisms to cell-size variations in mixer models can diverge to infinity. Plots of the noise contributions in (25) as a function of 1 + ‹*f*› (average increase in size in the timer phase) for two levels of 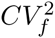 (timer-phase noise) when a timer precedes an adder (*a* = 1). Contributions are normalized by their value at ‹*f*› = 0. While the noise contribution of partitioning errors increases unboundedly with ‹*f*›, contributions from cell-division noise and growth-rate fluctuations first decrease and then increase. All noise contributions blow up at the same value of ‹*f*›, which decreases with increasing 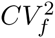

Ultimately all sensitivities in (25) diverge at the same value of ‹*f*› and this divergence corresponds to 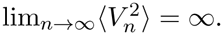. This phenomenon of size variance growing unbounded over time is a unique feature of mixer models absent in the generalized adder. The exact condition for the blow up can be traced back to the denominator of (23) becoming negative

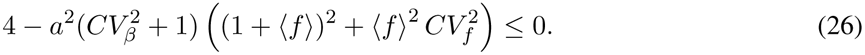

Since ‹*f*› < 1, (26) requires the presence of at least one noise mechanism 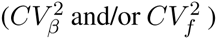 at sufficiently high levels. For example, in a timer-adder mixer model (*a* = 1), with a 75% increase in size in the timer phase (‹*f*› = 0.75), (26) holds when

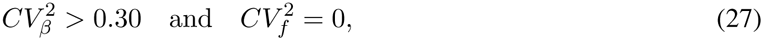

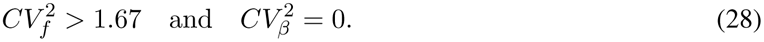

Note that the extent of partitioning errors required to destabilize the size variance is significantly smaller than the level of timer-phase noise. Finally, it is important to point out that the condition for variance divergence (26) depends on the timer-phase characteristics and partitioning errors, but is invariant of additive noise mechanism ∆_*n,α*_*n*__, and whether the timer follows or precedes the generalized adder.

## Power-law distribution of the newborn cell size

It turns out that the above result on infinite size variance can be generalized, in the sense that, for any non-zero noise levels 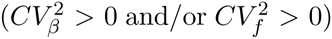 some higher-order moment of *V*_*n*_ grows unboundedly over time. This occurs in spite of all moments of iid random variables (*f*_*n*_, ∆_*n*_, *α*_*n*_) assumed to be finite. For example, consider the ideal case of zero partitioning errors, i.e., *β*_*n*_ = ‹*β*› with probability 1, and low timer-phase noise 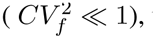 then

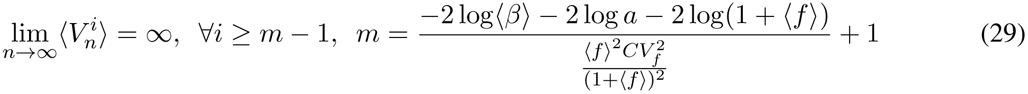

(SI section S3). Consistent with finding of [34], divergence of higher-order moments in (29) generates a power-law distribution for the newborn cell size. In particular, the steady-state distribution of *V*_*n*_ satisfies *p(x) ∝ x*^−*m*^ for sufficiently high values of *x* [35]. We refer to *m* as the power-law exponent arising form the mixer model structure, and it decreases with longer and noisier timer phase (increasing 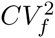 and ‹*f*›). Moreover, while *m* = *∞* for a sizer (*a* = 0), the exponent decreases as size control shifts towards the adder.

To investigate the effect of partitioning errors we model *β*_*n*_ via a beta distribution with mean *‹β›* and coefficient of variation squared 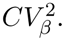. While analytical approximations for *m* are available in certain limits (see SI section S3), we numerically study *m* as a function of 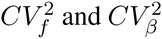 (Fig. 3). Our results shows that while the power-law exponent *m* decreases with increasing levels of either noise source, it is much more sensitive to partitioning errors. For example, 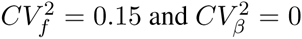 result in a power-law exponent of *m* = 60, but this value sharply drops to *m* = 15 in the opposite case of 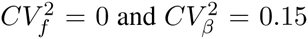 (Fig. 3). As observed in the previous section, the power-law exponent is independent of ∆_*n,α*_*n*__, and the order in which size control is executed. Next, we test these predictions with single-cell size measurements in *C. crescentus* that follows a mixer model for size homeostasis [32, 36].

**Figure 3:**
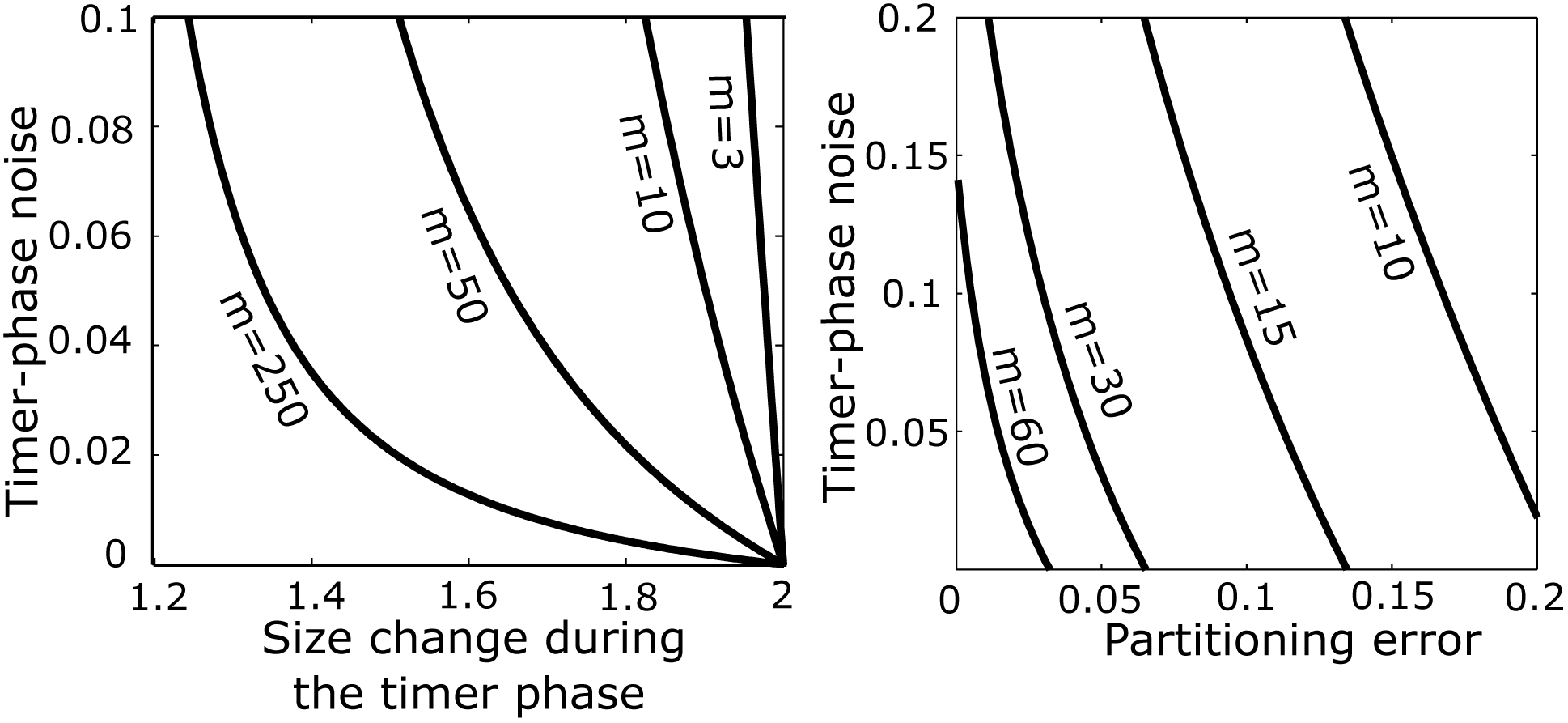
The power-law exponent of the newborn cell size distribution decreases with increasing levels of partitioning errors and noise in the timer phase. *Left*. Contour plot of the exponent *m* as a function of the mean size fold-change in the timer phase 1 + ‹*f*›, and the timer-phase noise 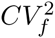, when partitioning errors are absent (*β*_*n*_ = 0.5 with probability 1). Each line corresponds to a constant value of *m*. Higher proportion of size increase occurring during the timer phase results in lower values of *m*, and hence lower-order moments of cell size grow unboundedly over time. *Right*. Plot of *m* as a function of the timer-phase noise (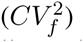) and partitioning errors (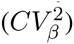) for ‹*f*› = 0.4 and *‹β›* = 0.5. Values of *m*are obtained numerically as described in SI section S3, and reveal that the exponent is more sensitive to partitioning errors. In particular, the value of *m* realized for a given level of partitioning error is much lower than its corresponding value for the same magnitude of timer-phase noise.

## Testing model predictions with *Caulobacter crescentus*

*C. crescentus* is a gram-negative bacterium where individual cells can exist in two forms – a mobile cell capable of swimming, and a “stalked” cell that has a tubular stalk enabling it to adhere to surfaces. Only the stalked cells are capable of cell division, and divide asymmetrically to produce a stalked, and a mobile cell, with the former being slightly larger in size. Size monitoring of individual stalked cells across multiple generations reveals a size homeostasis strategy based on a mixer model, where the timer follows a pure adder. More specifically, newborns grow exponentially for a size-independent time *t_n_* producing a 1 + *f*_*n*_ size fold-change in the timer phase with ‹*f*› ≈ 40% [32]. While there are considerable intercellular differences in *f*_*n*_, they are independent of the newborn size *V*_*n*_ [32]. This is consistent with our model assumption that in each cell cycle, *f*_*n*_ is drawn independently from a fixed distribution. The end of the timer phase marks the start of cell-wall constriction, and division occurs after adding a fixed size from the timer phase [32]. Similar to *f*_*n*_, the size added by individual cells is also observed to be independent of *V*_*n*_. Since in *C. crescentus*, growth rates exhibit considerable correlations across generations [32], independence of size added and *V*_*n*_ imply *c* = 0 in (8). This results in the following mixer model describing the stochastic dynamics of newborn size

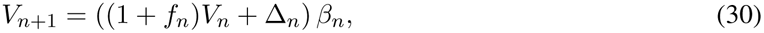

where *f*_*n*_, ∆_*n*_ and *β*_*n*_ are iid random variables describing stochasticity in the timer phase, size added during the adder phase, and the partitioning process, respectively. While *β*_*n*_ ∈ (0, 1) has finite moments by construction, all moments of *f*_*n*_, ∆_*n*_ are assumed to be bounded. We explore if the *C. crescentus* newborn size follows a power-law distribution with an exponent similar to as predicted by the mixer model.

Based on measurements of *V*_*n*_ we estimate significant noise in the timer phase 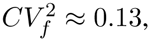, but precise partitioning of mother cell volume resulting in an order of magnitude lower partitioning errors 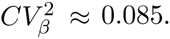. Since the experiment only tracks the stalked daughter cells that are slightly larger than the mobile daughter cells, we use ‹β› ≈ 0.58 as per measurements from [36]. We refer the reader to Table I in SI section S4 for precise parameter values with confidence intervals. Based on these parameters, the mixer model (30) predicts an exponent of *m* = 29 *±* 0.9, where the 95% confidence interval is obtained based on uncertainties in the estimates of underlying parameters (see SI section S4 for details). Intriguingly, consistent with the theoretical predictions, data reveals that the *C. crescentus* newborn cell size also follows

a power-law distribution (Fig. 4). However, the power-law exponent estimated from data using a maximum-likelihood method [35] is found to be *m*_*V*_ = 15.6 ± 1.1, and considerably smaller than *m* = 29 ± 0.9 as predicted by the mixer model structure.

**Fig. 4.**
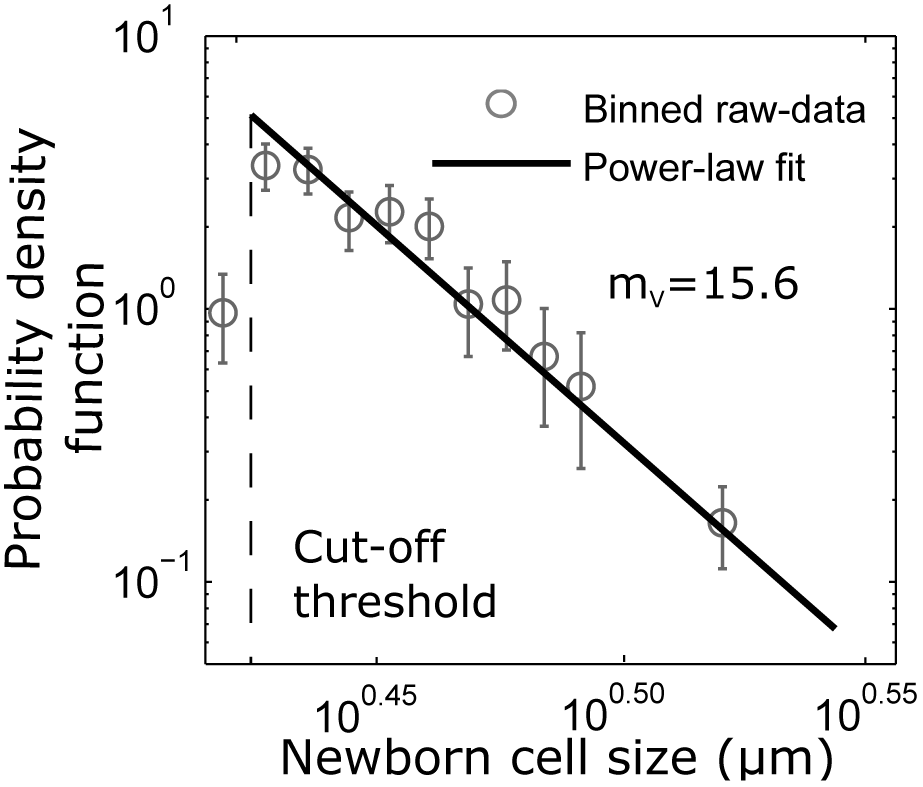
Consistent with model predictions, the *C. crescentus* newborn cell size follows a power-law distribution. The probability density function (pdf) is estimated by first dividing newborn sizes into different bins, and then finding the number of samples in each bin, divided by the product of the total number of samples and bin width. The error bars show the 95% confidence interval for bin height with the medians as the dots. After a size cutoff of 2.65 *µm* (≈ 460 cells), the pdf follows a power law distribution (linear in log-log scale). The power-law exponent *m_V_* = 15.6 1.1 ( denotes 95% confidence interval obtained using bootstrapping) is estimated using a maximum likelihood estimate method (SI section S5). The raw data is binned starting at 2.5861 *µm* using bin widths of 0.054 *µm* with the last bin 0.3779 *µm* wide.

Recall that the mixer model assumes all moments of ∆_*n*_ and *f*_*n*_ to be finite, and relaxing this assumption could explain the discrepancy between *m*_*V*_ and *m*. For example, consider the size fold-change 1 + *f*_*n*_ = exp(*α*_*n*_*t*_*n*_), where *α*_*n*_ is the growth rate, and *t*_*n*_ is the duration of the timer phase in the *n*^th^ cell cycle. If *α*_*n*_*t*_*n*_ is drawn independently from a Gaussian distribution, then size fold-change is lognormally distributed with finite moments. Since *α*_*n*_*t*_*n*_ is positively valued, perhaps a better approximation would be to draw *α*_*n*_*t*_*n*_ from a Gamma distribution, in which case, the size fold-change follows a power-law distribution. Thus, in principle, ∆_*n*_ and *f*_*n*_ could itself follow power-law distributions with exponents *m*_∆_ and *m*_*f*_. The exponent of the size distribution would then be essentially determined by the minimum of all the exponents *m*, *m*_∆_ and *m*_*f*_. To intuitively see this consider the size added ∆_*n*_ having infinite variance 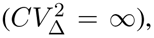, then as per (25) the steady-state size variance will also be infinite. Thus, *m*_*V*_ = *m*_∆_ = 3, irrespective of the value of *m*. Our data analysis shows that both ∆_*n*_ and *f*_*n*_ follow power-law distributions (see Fig. S1 in SI) with exponents *m*_∆_ = 13.5 *±* 1.9 and *m*_*f*_ = 19.6 *±* 2.3, respectively, that are smaller than *m* and much close to *m*_*V*_. In fact, the 95% confidence interval for the estimate of *m*_∆_ = 13.5 *±* 1.9 is overlapping with that of *m*_*V*_ = 15.6*±*1.1. In summary, while *C. crescentus* newborn sizes do indeed follow a power-law distribution as predicted by mixer model, their exponent is likely explained by the power-law statistics of the size added by single cells in the adder phase.

## Discussion

Diverse cell types employ different size-control strategies to maintain an optimal cell size. How effective are these strategies in regulating stochastic variation in cell size that arises from various physiologically relevant noise sources? We addressed this question in the context of the generalized added, a recently uncovered size-homeostasis mechanism in microbes, where timing of cell division is regulated such that successive newborn cell sizes are related via (9). In this framework, *a* = 1 corresponds to a pure adder (division occurs after adding size ∆_*n,α*_*n*__ from birth), and *a* = 0, a sizer (division occurs when size reaches ∆_*n,α*_*n*__). A key and novel assumption in our model is the specific form for ∆_*n,α*_*n*__ that is motivated by experimental findings [14]. In particular, ∆_*n,α*_*n*__ was assumed to be a product of an iid random variable ∆_*n*_ with finite moments that is drawn independently of size, and a (linear or exponential) function of the cellular growth rate (7).

Our main result for the generalized adder connects the stochastic variation in cell size 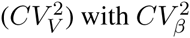 (errors in the partitioning process), 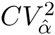 (growth-rate fluctuations) and 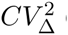 (noise in ∆_*n*_ or cell-division noise). In the limit of low noise, 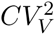 can be decomposed as a linear function of the different noise sources (16), with the coefficients representing the sensitivity of 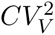 to the corresponding noise source. This formula reveals that for 0 *< a ≤* 1, the size variation is most sensitive to partitioning errors, and not surprisingly this process in highly regulated in many microbes. Cell division in bacteria is mediated by the septal ring, and spatial precision in ring formation essentially dictates the partitioning error. In *E. coli*, the positioning of the septal ring at the cell midpoint is actively regulated using the Min protein system [12, 13], and mutations in the Min proteins can significantly amplify cell-to-cell size variations due to large partitioning errors [22]. Intriguingly, while the sensitivity of 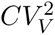 to 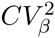 increases with *a*, sensitivities to other noise sources (growth-rate fluctuations and cell-division noise) decreases with *a* (Fig. 1). This implies that the adder (*a* = 1) is effective in minimizing 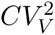 from the additive noise source ∆_*n,α*_*n*__, but it is susceptible to multiplicative noise arising through the partitioning process. In comparison, the sizer (*a* = 0) performs better in buffering size variations from partitioning errors (Fig. 1). Overall, this analysis suggests that organisms with asymmetric partitioning (large 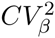 values), should employ sizer-based control to regulate variations around an optimal size. On the other hand, the adder provides better suppression of stochastic size deviations in organisms with symmetric and highly regulated partitioning (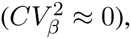), where cell-division noise or growth-rate fluctuations may be the dominant noise mechanisms. Finally, given that sensitivities in (16) have different functional dependencies on *a*, it is easy to generate scenarios where 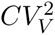 is minimized at an intermediate value of *a* when multiple noise sources are present (Fig. 1).

We further expanded the size computations to mixer models of size control, where a timer phase precedes or follows a generalized adder. It is well known that a simple timer mechanism for controlling cell-division does not provide size homeostasis, in the sense that, the variance in cell size grows unboundedly over time [37]. Our results show that mixing a timer with an generalized adder results in similar instabilities - all moments of the newborn cell size of order *m −* 1 and higher grow unboundedly over time, where *m* is given by (29). This leads to the cell size following a power-law distribution with exponent *m* [34], which decreases with increasing *a* (size control shifts towards the adder), and for increasing stochasticity in the partitioning process and timer phase (Fig. 3). Interestingly, when partitioning errors are negligible, adding a timer phase can reduce 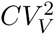 (Fig. 2). This leads to a counter-intuitive result that in some cases, mixer models may have lower stochastic variations in cell size as compared to the generalized adder, even though higher-order moments diverge in the mixer but not in the latter. Moreover, we also find that the generalized adder followed by a timer is more noisy than the reverse scenario (see (24)), but both systems exhibit the same power-law exponent *m*.

The prediction of power-law distribution in mixer models was tested with single-cell data on *C. crescentus*, where size regulation is mediated through a timer followed by an adder [36]. Consistent with theory, we find strong evidence of the newborn cell size following a power-law distribution with an exponent of *m*_*V*_ ≈ 15.6 (Fig. 4). Surprisingly, independent measurements of noise sources and cell-cycle parameters from [32, 36], predicted an exponent two-fold higher as per (29). We provide a simple mechanistic explanation for this discrepancy that lies in statistical fluctuations in *f*_*n*_ (size fold change in timer phase) or ∆_*n*_ (size added during adder phase) itself following power-law distributions, i.e., their moments diverge beyond a certain order. It turn out that in *C. crescentus*, the power-law exponents of *f*_*n*_ and ∆_*n*_ are much smaller than as predicted by the mixer-model structure, and hence dominate in terms of determining the power-law exponent of the size distribution (Fig. S1).

In summary, this study provides a systematic understanding of how stochastic variations in cell size are regulated in the context of different size homeostasis strategies, and different underlying noise mechanisms. Inspired from data [14, 22], we took a phenomenological approach to capturing noise in the cell-cycle process and growth-rate fluctuations through ∆_*n,α*_*n*__. However, future work will consider mechanistic ways of incorporating noise by explicitly modeling the cell cycle, as in [38–43]. Moreover, by linking cell size to expression of constitutive genes we hope to uncover strategies for maintaining concentrations of essential proteins [44–49], in spite of stochastic and temporal changes in cell size.

## SUPPLEMENTARY MATERIAL

An online supplement to this article can be found by visiting bioRxiv online at http://biorxiv.org/.

## Acknowledgments

AS is supported by the National Science Foundation Grant DMS-1312926.

## References

1. Alison C. Lloyd. The Regulation of Cell Size. Cell, 154:1194–1205, 2013.

2. Arieh Zaritsky and Conrad L. Woldringh. Chromosome replication, cell growth, division and shape: a personal perspective. Microbial Physiology and Metabolism, 6:756, 2015.

3. Arieh Zaritsky. Cell-Shape Homeostasis in Escherichia coli Is Driven by Growth, Division, and Nucleoid Complexity. Biophysical Journal, 109:178–181, 2015.

4. Ping Wang, Lydia Robert, James Pelletier, Wei Lien Dang, Francois Taddei, Andrew Wright, and Suckjoon Jun. Robust growth of escherichia coli. Current biology, 20:1099–1103, 2010.

5. Wallace F. Marshall, Kevin D. Young, Matthew Swaffer, Elizabeth Wood, Paul Nurse, Akatsuki Kimura, Joseph Frankel, John Wallingford, Virginia Walbot, Xian Qu, and Adrienne HK Roeder. What determines cell size? BMC Biology, 10:101, 2012.

6. Michael Halter, John T. Elliott, Joseph B. Hubbard, Alessandro Tona, and Anne L. Plant. Cell volume distributions reveal cell growth rates and division times. Journal of Theoretical Biology, 257:124–130, 2009.

7. Amit Tzur, Ran Kafri, Valerie S. LeBleu, Galit Lahav, and Marc W. Kirschner. Cell Growth and Size Homeostasis in Proliferating Animal Cells. Science, 325:167–171, 2009.

8. Yu Tanouchi, Anand Pai, Heungwon Park, Shuqiang Huang, Rumen Stamatov, Nicolas E. Buchler, and Lingchong You. A noisy linear map underlies oscillations in cell size and gene expression in bacteria. Nature, 523:357–360, 2015.

9. Lydia Robert, Marc Hoffmann, Nathalie Krell, Stéphane Aymerich, Jérôme Robert, and Marie Doumic. Division in Escherichia coli is triggered by a size-sensing rather than a timing mechanism. BMC Biology, 12:17, 2014.

10. H. E. Kubitschek. Growth during the bacterial cell cycle. analysis of cell size distribution. Biophysical Journal, 9:792–809, 1969.

11. An-Chun Chien, Norbert S Hill, and Petra Anne Levin. Cell size control in bacteria. Current Biology, 22:R340–R349, 2012.

12. A. G. Marr, R. J. Harvey, and W. C. Trentini. Growth and division of escherichia coli. Journal of Bacteriology, 91:2388–2389, 1966.

13. Jaan Mannik, Fabai Wu, Felix Hol, Paola Bisicchia, David Sherratt, Juan Keymer, and Cees Dekker. Robustness and accuracy of cell division in escherichia coli in diverse cell shapes. Proceedings of the National Academy of Sciences, 109:6957–6962, 2012.

14. Sattar Taheri-Araghi, Serena Bradde, John T. Sauls, Norbert S. Hill, Petra Anne Levin, Johan Paulsson, Massimo Vergassola, and Suckjoon Jun. Cell-size control and homeostasis in bacteria. Current Biology, 25:385–391, 2015.

15. Daniel J. Kiviet, Philippe Nghe, Noreen Walker, Sarah Boulineau, Vanda Sunderlikova, and Sander J. Tans. Stochasticity of metabolism and growth at the single-cell level. Nature, 514:376–379, 2014.

16. Francisco Ferrezuelo, Neus Colomina, Alida Palmisano, Eloi Gar, Carme Gallego, Attila Csiksz-Nagy, and Mart Aldea. The critical size is set at a single-cell level by growth rate to attain homeostasis and adaptation. Nature communications, 3:1012, 2012.

17. Mikihiro Hashimoto, Takashi Nozoe, Hidenori Nakaoka, Reiko Okura, Sayo Akiyoshi, Kunihiko Kaneko, Edo Kussell, and Yuichi Wakamoto. Noise-driven growth rate gain in clonal cellular populations. Proceedings of the National Academy of Sciences, 113:3251–3256, 2016.

18. Stephen Vadia and Petra Anne Levin. Growth rate and cell size: a re-examination of the growth law. Current Opinion in Microbiology, 24:96–103, 2015.

19. Lydia Robert. Size sensors in bacteria, cell cycle control, and size control. Frontiers in Microbiology, 6:515, 2015.

20. Akos Sveiczer, John J. Tyson, and Bela Novak. A stochastic, molecular model of the fission yeast cell cycle: role of the nucleocytoplasmic ratio in cycle time regulation. Biophysical Chemistry, 92:1–15, 2001.

21. Stefano Di Talia, Jan M. Skotheim, James M. Bean, Eric D. Siggia, and Frederick R. Cross. The effects of molecular noise and size control on variability in the budding yeast cell cycle. Nature, 448:947–951, 2007.

22. Manuel Campos, Ivan V. Surovtsev, Setsu Kato, Ahmad Paintdakhi, Bruno Beltran, Sarah E. Ebmeier, and Christine Jacobs-Wagner. A constant size extension drives bacterial cell size homeostasis. Cell, 159:1433–1446, 2014.

23. Ariel Amir. Cell size regulation in bacteria. Physical Review Letters, 112:208102, 2014.

24. John T Sauls, Dongyang Li, and Suckjoon Jun. Adder and a coarse-grained approach to cell size homeostasis in bacteria. Current Opinion in Cell Biology, 38:38–44, 2016.

25. Ilya Soifer, Lydia Robert, and Ariel Amir. Single-cell analysis of growth in budding yeast and bacteria reveals a common size regulation strategy. Current Biology, 26:356–361, 2016.

26. Maxime Deforet, Dave van Ditmarsch, and João B. Xavier. Cell-size homeostasis and the incremental rule in a bacterial pathogen. Biophysical Journal, 109:521–528, 2015.

27. Anouchka Fievet, Adrien Ducret, Tâm Mignot, Odile Valette, Lydia Robert, Romain Pardoux, Alain Roger Dolla, and Corinne Aubert. Single-cell analysis of growth and cell division of the anaerobe Desulfovibrio vulgaris Hildenborough. Frontiers in Microbiology, 6:1378, 2015.

28. Suckjoon Jun and Sattar Taheri-Araghi. Cell-size maintenance: universal strategy revealed. Trends in Microbiology, 23:4–6, 2015.

29. Kally Z. Pan, Timothy E. Saunders, Ignacio Flor-Parra, Martin Howard, and Fred Chang. Cortical regulation of cell size by a sizer cdr2p. eLife, 3:e02040, 2014.

30. Jonathan J. Turner, Jennifer C. Ewald, and Jan M. Skotheim. Cell size control in yeast. Current Biology, 22:R350–R359, 2012.

31. Matteo Osella, Eileen Nugent, and Marco Cosentino Lagomarsino. Concerted control of Escherichia coli cell division. Proceedings of the National Academy of Sciences, 111:3431–3435, 2014.

32. Shiladitya Banerjee, Klevin Lo, Thomas Kuntz, Matthew K. Daddysman, Aaron R. Dinner, and Norbert F. Scherer. Crossover in the dynamics of cell wall growth controls bacterial division times. bioRxiv 047589, 2016. http://biorxiv.org/content/early/2016/04/07/047589.

33. Frederick R. Cross and James G. Umen. The Chlamydomonas cell cycle. The Plant Journal: For Cell and Molecular Biology, 82:370–392, 2015.

34. Andrew Marantan and Ariel Amir. Stochastic modeling of cell growth with symmetric or asymmetric division. Physical Review E, 94:012405, 2016.

35. Aaron Clauset, Cosma Rohilla Shalizi, and M.E.J. Newman. Power-law distributions in emperical data. SIAM Review, 51:661–703, 2009.

36. Srividya Iyer-Biswas, Charles S Wright, Jonathan T Henry, Klevin Lo, Stanislav Burov, Yihan Lin, Gavin E Crooks, Sean Crosson, Aaron R Dinner, and Norbert F Scherer. Scaling laws governing stochastic growth and division of single bacterial cells. Proceedings of the National Academy of Sciences, 111:15912–15917, 2014.

37. Cesar Augusto Vargas-Garcia, Mohammad Soltani, and Abhyudai Singh. Conditions for cell size homeostasis: A stochastic hybrid systems approach. arXiv:1606.00535v1 [q-bio.CB], 2016. https://arxiv.org/abs/1606.00535.

38. Khem Raj Ghusinga, Cesar A. Vargas-Garcia, and Abhyudai Singh. A mechanistic stochastic framework for regulating bacterial cell division. Scientific Reports, 6:30229, 2016.

39. Leigh K. Harris and Julie A. Theriot. Relative rates of surface and volume synthesis set bacterial cell size. Cell, 165:1479–1492, 2016.

40. Mats Wallden, David Fange, Ebba Gregorsson Lundius, zden Baltekin, and Johan Elf. The synchronization of replication and division cycles in individual e. coli cells. Cell, 166:729–739, 2016.

41. Bela Novak and John J. Tyson. Modeling the cell division cycle: M-phase trigger, oscillations, and size control. Journal of Theoretical Biology, 165:101–134, 1993.

42. Duarte Antunes and Abhyudai Singh. Quantifying gene expression variability arising from randomness in cell division times. Journal of Mathematical Biology, 71:437–463, 2014.

43. Vahid Shahrezaei and Samuel Marguerat. Connecting growth with gene expression: of noise and numbers. Current Opinion in Microbiology, 25:127–135, 2015.

44. Olivia Padovan-Merhar, Gautham P. Nair, Andrew G. Biaesch, Andreas Mayer, Steven Scarfone, Shawn W. Foley, Angela R. Wu, L. Stirling Churchman, Abhyudai Singh, and Arjun Raj. Single mammalian cells compensate for differences in cellular volume and DNA copy number through independent global transcriptional mechanisms. Molecular Cell, 58:339–352, 2015.

45. Mohammad Soltani, Cesar Vargas, Duarte Antunes, and Abhyudai Singh. Intercellular variability in protein levels from stochastic expression and noisy cell cycle processes. PLoS Computational Biology, 12:e1004972, 2016.

46. Mohammad Soltani and Abhyudai Singh. Cell-cycle coupled expression minimizes random fluctuations in gene product levels. arXiv:1605.02251 [q-bio], 2016. http://biorxiv.org/content/biorxiv/early/2016/05/08/052159.

47. Noreen Walker, Philippe Nghe, and Sander J. Tans. Generation and filtering of gene expression noise by the bacterial cell cycle. BMC Biology, 14:1–10, 2016.

48. Hermannus Kempe, Anne Shwabe, Frederic Cremazy, Pernette J. Verschure, and Frank J. Bruggeman. The volumes and transcript counts of single cells reveal concentration homeostasis and capture biological noise. Molecular Biology of the Cell, 26:797–804, 2014.

49. Kurt M. Schmoller and Jan M. Skotheim. The Biosynthetic Basis of Cell Size Control. Trends in Cell Biology, 25:793–802, 2015.

